# 3’UTR variants of ALS-linked RNAs modify subcellular and cellular phenotypes

**DOI:** 10.1101/2025.04.05.647382

**Authors:** Melis Savasan-Sogut, Danae Campos-Melo, Michael J. Strong

**Author notes:** Corresponding author: (DCM), (MJS). These authors contributed equally to the work (Melis Savasan-Sogut, Danae Campos-Melo).

## Abstract

While most human genes express mRNA 3’untranslated region (3’UTR) variants of different lengths, their impact on cell physiology and disease remains largely unknown. Here, we studied 3’UTR length heterogeneity in amyotrophic lateral sclerosis (ALS) and determined that three ALS-linked transcripts exhibit lengthening of their 3’UTRs in patient samples. We investigated phenotypical effects in a neuronal cell line expressing these 3’UTRs and observed that expression of these unique 3’UTRs induces morphological changes at different levels. Among the most expressed 3’UTRs variants in ALS, *NEFH* 3’UTR-Long induces the formation of nuclear RNA clusters and *SOD1* 3’UTR-Long diminishes filopodia in the plasma membrane. *SQSTM1* 3’UTR-Long did not show major changes in nuclear RNA clusters or filopodia. This is the first report that suggests that 3’UTRs may function independent of the coding region and modify the phenotype of a cell, further expanding the impact of alterations in mRNA biogenesis in ALS.

## Introduction

MRNA 3’ untranslated regions (UTRs) are sequences located at the 3’end of the protein-coding region of mRNAs. Variants of 3’UTRs are naturally generated for approximately 70% of human genes through different processes including alternative cleavage and polyadenylation (APA), which results in transcripts that encode for the same protein but differ in the length of their 3’ untranslated sequences. Cleavage and polyadenylation occur co-transcriptionally and are executed by several multisubunit protein complexes, including poly(A) polymerase (PAP) and associated factors. 3’UTR variants are expressed in a gene and cell-type-specific manner, containing distinct *cis*-regulatory elements such as binding sites for RNA-binding proteins and miRNAs, in addition to expressing *Alu*, AU- and GU-rich elements (1–3).

3’UTRs have roles at different stages of the mRNA life cycle, such as RNA stability, localization and translation efficiency (4). They can also function as scaffolds for protein binding to nascent proteins during translation to direct their transport and function (5). APA was initially thought to be a gene expression regulatory mechanism but recently it was observed that mRNA amounts and 3’UTR lengths are independent processes that determine levels and spatial organization of protein synthesis (6). Moreover, widespread differential expression of cognate coding sequences (CDS) and 3’UTRs have been reported in different species in the nervous system and other tissues, suggesting additional functions for 3’UTRs independent of their cognate coding sequences (7–9).

The association between mRNA 3’UTRs and diseases has recently gained increasing attention, especially in cancer biology (10). However, the function of 3’UTR variants in other diseases is largely unknown. In amyotrophic lateral sclerosis (ALS), our group and others have described alterations in the RNA metabolism at multiple levels (11–13). Among them, APA defects such as increased use of proximal polyadenylation sites in transcripts in the cerebellum of C9ORF72 patients have been reported (14). More recently, variants of the interleukin 18 receptor accessory protein 3′UTR that protect against ALS have been described (15).

In this study, we explored APA changes in the spinal cord of ALS and control patients. We performed RNA-seq and quantitative alternative polyadenylation analysis (QAPA) (16) and observed dysregulation of 3’UTR variants in a group of transcripts, including lengthening of mRNA 3’UTRs of RNAs whose expression has been shown to be altered in ALS. We cloned and expressed EGFP-linked 3’UTR variants of *NEFH*, *SOD1,* and *SQSTM1* transcripts in neuronal cells and studied changes at the subcellular and cellular levels induced by these 3’UTRs using single-molecule fluorescence *in situ* hybridization (smFISH). We observed that in the absence of coding sequences, 3’UTR variants of the same transcript not only exhibit different patterns of localization within the cell but also determine specific characteristics. Among the 3’UTR variants that show lengthening in ALS, *NEFH* 3’UTR-Long increases the formation of nuclear RNA clusters and *SOD1* 3’UTR-Long reduces the formation of filopodia. *SQSTM1* 3’UTR-Long did not show difference in RNA cluster and filopodia formation. Together, our results suggest that mRNA 3’UTR sequences can directly influence the phenotype of the cell and might contribute to the disease pathology.

## Results

We performed RNA-seq and QAPA analysis of human ALS and control spinal cord tissues. We observed that 5.6% of the 3’UTRs were dysregulated in the disease (Fig 1A and S1 Fig). In this group, 1,169 and 1,195 3’UTR variants were down and upregulated in ALS, respectively. The distribution of DPAU data exhibited a moderate shift in APA preference for the majority of down- and upregulated-3’UTRs (Fig 1B). Reactome enrichment of the 3’UTR dysregulation in ALS showed pathways involved in cellular responses to stress and membrane trafficking, which have been associated with the disease (Fig 1C) (17,18). Reactome enrichment of 3’UTR lengthening highlighted several processes also associated with ALS, such as endo-lysosomal pathway, splicing, autophagy and ferroptosis (19–22)(Fig 1D). Among those 3’UTR variants for which expression was altered, we found a group of ALS-linked transcripts (Table 1). The majority have functions in the cellular pathways/processes shown in the reactome enrichment. Some of these mRNAs express a small number of 3’UTRs of different lengths (e.g. *NEFH*), while others express many (e.g. *TARDBP*). Three of the ALS-associated transcripts, *NEFH*, *SOD1,* and *SQSTM1,* showed lengthening of their 3’UTRs and were selected for further experiments.

**Figure 1.**
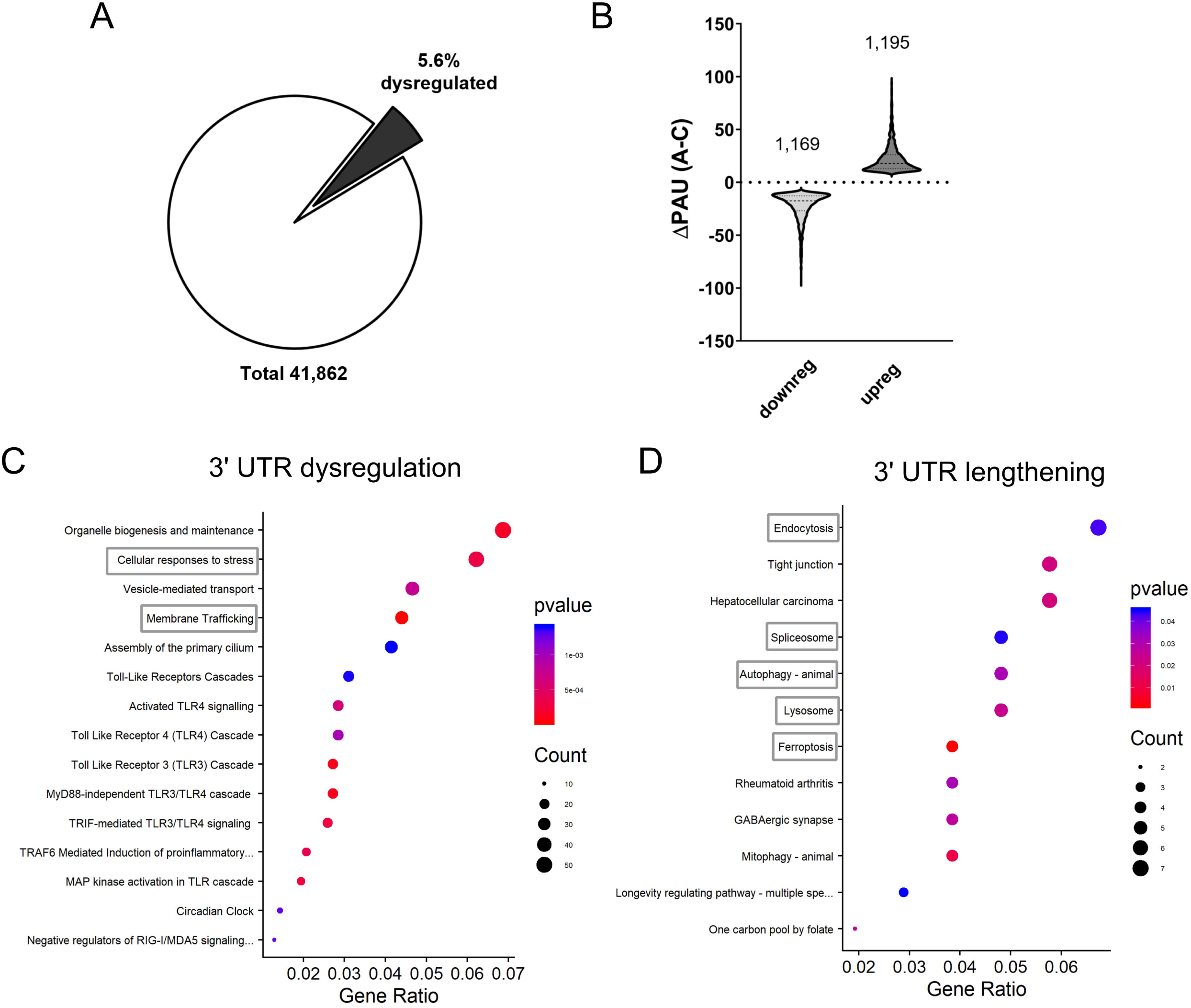
QAPA of RNA-seq from ALS spinal cords. Quantitative alternative polyadenylation analysis (QAPA) was used to study changes in the polyadenylation site usage (PAS) in ALS relative to all mRNA 3′UTR isoforms of each gene. (A) A significant group of 3’UTRs are dysregulated in ALS. (B) The change in poly(A) site usage (DPAU) between ALS (A) and control (C) shows data distribution for down and upregulated 3’UTRs. Numbers above the violin plots indicate the number of 3’UTRs down or upregulated. (C) Reactome enrichment of 3’UTR dysregulation in ALS. (D) KEGG enrichment of 3’UTR lengthening in ALS. Statistically significant pathways altered in ALS are listed in (C) and (D). Circle sizes indicate the UTR count and colors the p-values. Dotted boxes in (C) and (D) indicate cellular processes previously reported as altered in ALS.

**Table 1.**
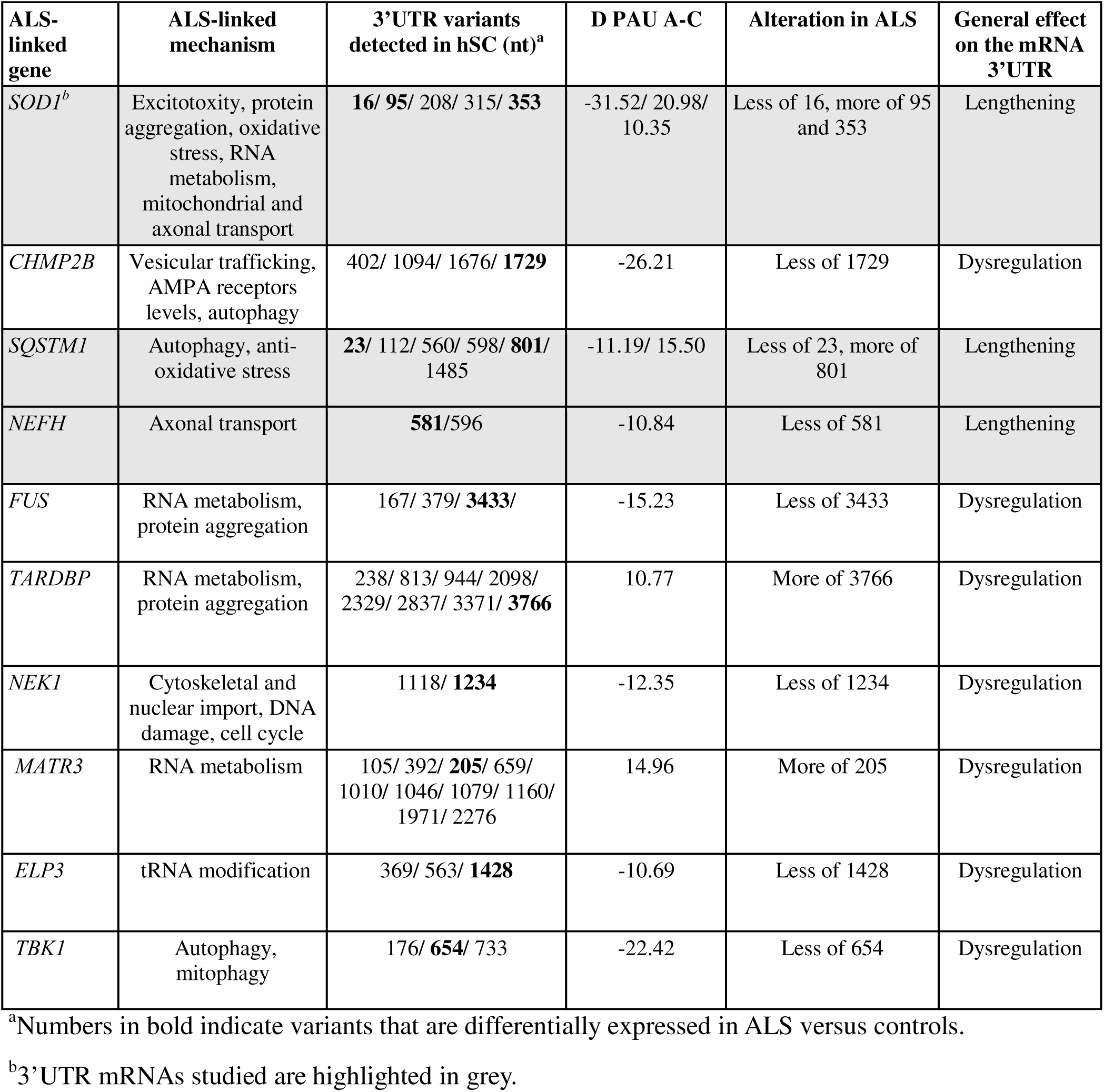
3’UTRs dysregulation in ALS-linked transcripts.

We expressed different variants of *NEFH*, *SOD1,* and *SQSTM1* 3’UTRs in a neuronal cell line (Fig 2A and 3A, S2A). For the quantification of different structures, only GFP+ cells expressing low levels of the fluorescent protein and showing healthy nuclei were considered. Confocal images of *NEFH* 3’UTR localization showed differences in the number of RNA clusters for the distinct variants in SH-SY5Y cells (Fig 2B). We quantified these *NEFH* 3’UTR clusters in the nucleus and the cytoplasm, and classified them into two size groups, small (<1.3 µm²) and large (>1.3 µm²). We determined that *NEFH* 3’UTR-Long formed a higher number of nuclear clusters than either *NEFH* 3’UTR-Short or control (Short: p=0.0017, control: 0.0252; Figure 2C). We observed an increased cluster size in the *NEFH* 3’UTR-Long group compared to the *NEFH* 3’UTR-Short variant (p=0.0170; Fig 2D). In addition, we observed that *NEFH* 3’UTR-Short formed a higher number and size of cytoplasmic clusters than the NEFH 3’UTR-Long variant (number of clusters p<0.0001; area of clusters p = 0.0004; Fig 2C and 2D, S3). *SQSTM1* 3’UTR variants did not show major differences in cluster and filopodia formation (S2 Fig A, B and C).

**Figure 2.**
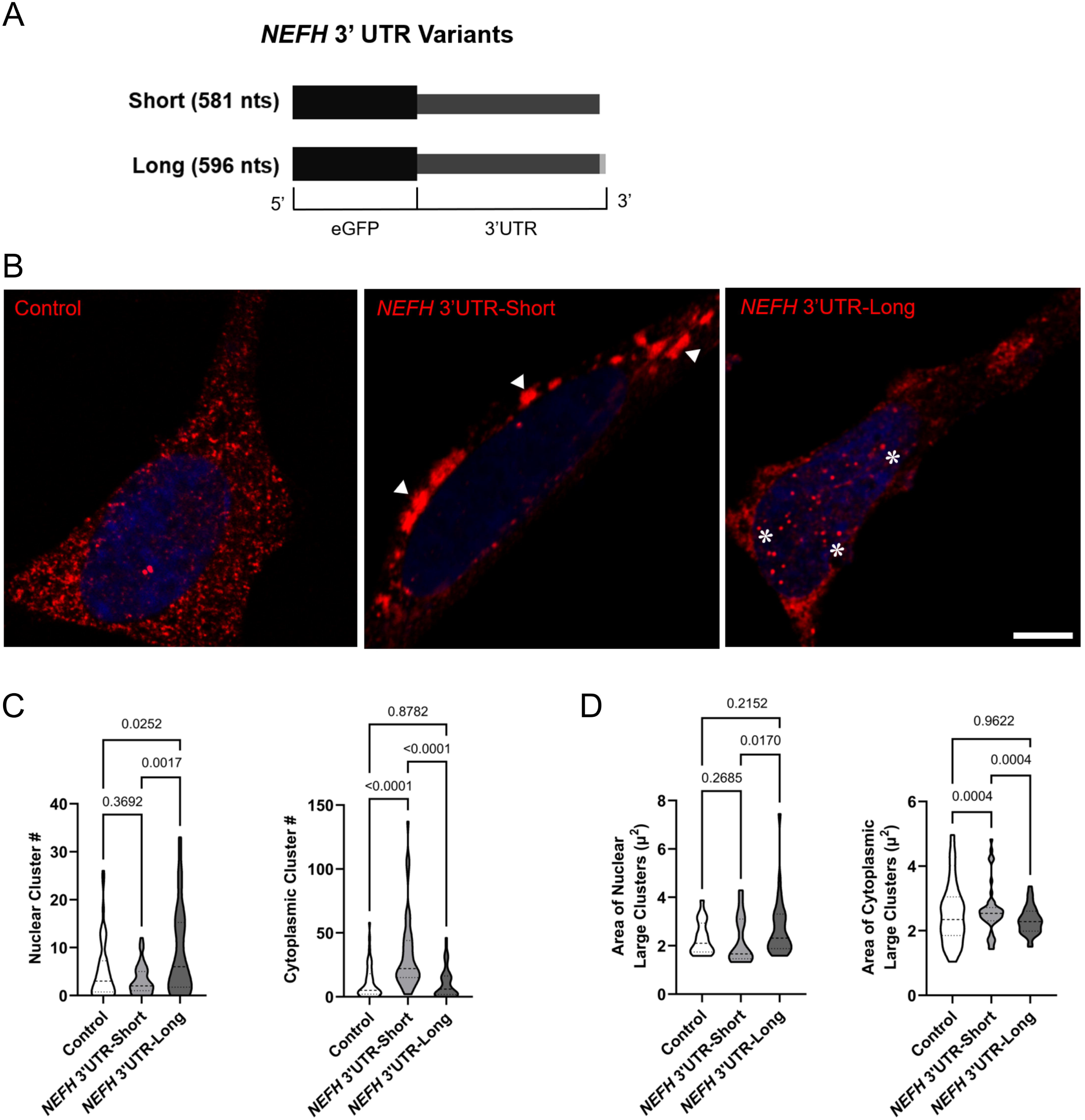
*NEFH* 3′UTR variants differentially induce RNA cluster formation. (A) Schematic representation of the *NEFH* 3′UTR variants. The 15-nt difference between the two variants located at the 3′ end of the long variant is indicated. (B) Subcellular localization of *NEFH* 3′UTR variants using RNA-FISH in differentiated SH-SY5Y cells transfected with eGFP-alone (Control), *NEFH* 3′UTR-Short, or Long (red). Arrowheads show cytoplasmic RNA clusters of 3′UTR-Short variant, whereas asterisks indicate nuclear clusters of 3′UTR-Long. Scale bar, 5 μm. (C) Quantification of clusters of *NEFH* 3′UTR variants. Violin plots show the number of RNA clusters in the nucleus and cytoplasm for Control, *NEFH* 3′UTR-Short, and Long. (D) Quantification of cluster areas of *NEFH* 3′UTR variants. Violin plots of RNA clusters areas in the nucleus and cytoplasm in Control, *NEFH* 3′UTR-Short, and Long. Statistical comparisons were conducted using a nonparametric Kruskal–Wallis test followed by original FDR correction. P-values are indicated above each comparison.

**Figure 3.**
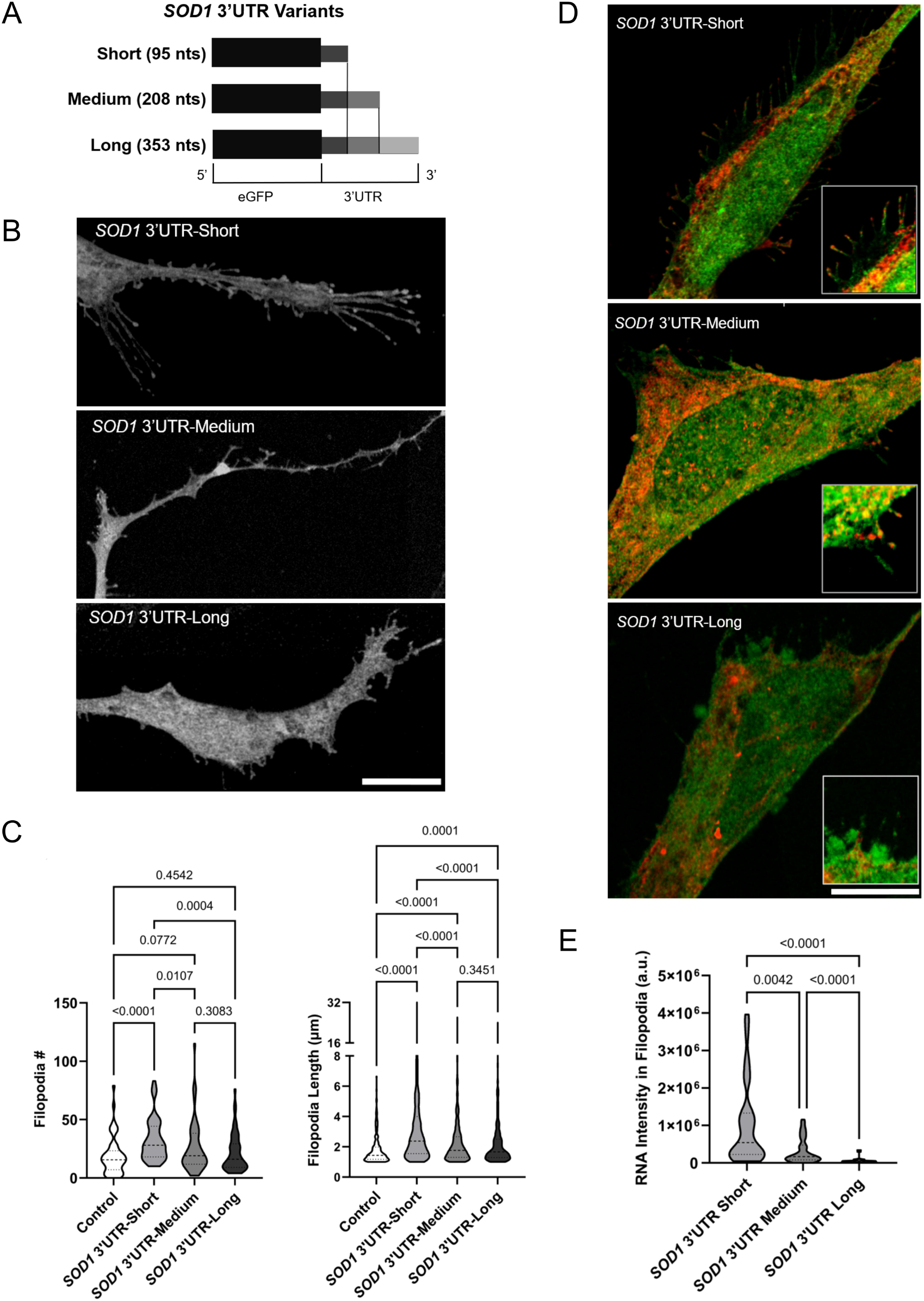
*SOD1* 3′UTR variants differentially induce filopodia formation. (A) Schematic representation of the *SOD1* 3′UTR variants. (B) Effect of *SOD1* 3’UTR variants on filopodia formation in differentiated SH-SY5Y cells transfected with *SOD1* 3′UTR-Short, Medium, or Long constructs. Scale bar, 5 μm. (C) Quantification of filopodia. Violin plots show the quantification of filopodia number and filopodia length for each group. (D) SOD1 3’UTRs in filopodia. Insets of pictures show a magnified view of the presence of *SOD1* 3’UTR variants at the distal tip of filopodia. Scale bar, 5 μm. (E) Quantification of SOD1 3’UTRs in filopodia. Violin plots represent RNA fluorescence intensity for each *SOD1* 3′UTR variant in filopodia.

Next, we studied changes in the morphology of the cells when *SOD1* 3’UTR variants were expressed (Fig 3A). We specifically focused on filopodia, thin finger-like protrusions of the plasma membrane that have variable length and mechanosensing, guidance and communication functions (23,24). Confocal images showed that *SOD1* 3′UTR variants differed in the filopodia abundance (Fig 3B). Quantification showed that cells expressing *SOD1* 3’UTR-Long have a significant reduction in the number and length of filopodia compared with cells expressing *SOD1* 3’UTR-Short (number of filopodia, Short: p= 0.0004; length of filopodia, Short and Control, p<0.0001; Fig 3C).

To determine whether filopodia changes were related to the localization of 3’UTR variants in these structures, we measured the amount of *SOD1* 3’UTR RNA (Fig 3D). The amount of *SOD1* 3′UTR-Long was significantly lower in filopodia than in Short and Medium 3′UTR variants (p<0.0001; Fig 3E). Quantification of lamellipodia, sheet-like protrusions that often coexist with filopodia in cells and also have mechanosensing functions (25), showed that *SOD1* 3′UTR-Long expression led to a trend in the reduction of the number of lamellipodia compared to Short variant (p=0.0848; S4 Fig). Finally, we examined the colocalization of *SOD1* 3’UTR variants with F-actin, the building blocks of filopodia and lamellipodia. We observed that *SOD1* 3’UTR-Medium and Long form clusters in the cytoplasm with F-actin while *SOD1* 3′UTR-Short distributed throughout the compartment. The *SOD1* 3’UTR-Medium variant was also present in clusters with F-actin in distal neurites, and the Short variant showed the tendency to localize near actin-rich protrusions (S5 Fig).

Altogether, our data suggest that 3’UTR expression is significantly altered in ALS, including ALS-linked transcripts. MRNA 3’UTRs that showed lengthening, *NEFH*, *SQSTM1* and *SOD1*, can alter the subcellular (RNA clusters) and the cellular (filopodia/lamellipodia) phenotypes in neuronal cells, in a pathological-like manner.

## Discussion

In this study, we observed that the expression of a group of 3’UTRs is altered in ALS and that these changes are associated with malfunctions in the disease. Among the dysregulated mRNA 3’UTRs, we found ALS-linked transcripts, such as *NEFL*, *SOD1,* and *SQSTM1*, that showed lengthening of their sequences. The expression of these mRNA 3’UTRs in a neuronal cell line showed that among the most expressed 3’UTRs variants in ALS, *NEFH* 3’UTR-Long and *SOD1* 3’UTR-Long, augment RNA clusters in the nucleus and diminish the number and length of filopodia in the membrane, respectively. *SQSTM1* 3’UTR-Long exhibited no alterations in nuclear RNA clusters or filopodia. These results suggest for the first time that 3’UTRs may function independently of coding regions and modify the phenotype of a cell.

The roles of mRNA 3’UTRs are not fully understood. Even though these sequences have been associated with post-transcriptional processes of the RNA, the choice of APA sites has lately been shown to have minimal effects on stability and translational efficiency (26,27). The lack of knowledge is partly due to the limited studies that explore in detail the functions of 3’UTRs without their cognate coding regions.

Mercer and col. described the expression of distinct 3’UTR-derived RNAs (7). Then, widespread independent expression of mRNA 3’UTRs in neurons and other cells was reported, including a broad range of 3’UTR-to-CDS expression ratios in different cell types that are spatially graded or change with developmental age (8). In human and mouse brain aging, there is an accumulation of isolated ribosome-associated 3′UTRs that tightly correlates with oxidative stress (28). Depending on the gene, isolated 3’UTRs might be generated by *de novo* transcription or post-transcriptional cleavage (9,29,30). Among the few studies that investigated the independent functions of 3’UTRs, it has been shown that 3’UTRs can be regulators that suppress tumor formation (31,32) and contain sequences that work as sensors of stress and changes in the cell *milieu*, such as growth factor levels (33).

Here, we showed that the expression of the 3’UTR-Long variant of *NEFH* induces the formation of RNA clusters in the nucleus. Mechanisms of RNA clustering have been studied in different models, such as mRNAs of germ granules in *Drosophila* (34), and it has also been associated with pathological states. RNA foci have been broadly involved in repeat-expansion disorders, like fragile X-associated tremor ataxia syndrome (FXTAS) and ALS (35,36). In C9ALS/FTD patients, either GGGGCC or CCCCGG repeats with atypical secondary structures are thought to form foci and interact with RNA-binding proteins, triggering a molecular cascade that leads to neurodegeneration (36). Consequently, the formation of nuclear clusters could induce retention and regulation of RNA expression and function.

Our observations that filopodia decrease when *SOD1* 3’UTR-Long is expressed and the formation of cytoplasmic clusters with F-actin might indicate that actin nucleation, crucial for filopodia/lamellipodia formation (37), is being blocked. The direct or indirect participation of *SOD1* 3’UTR-Long in this process and, ultimately, in the sensing and communication between cells is an interesting focus of research. Further studies are necessary to elucidate these ideas and the contribution of these mechanisms to the pathogenesis of ALS.

Non-coding RNAs are diverse in structure and function. MRNAs support versatility as 3’UTRs can act as part of a larger coding molecule conferring it specific characteristics or in an independent manner may be similar to long noncoding RNAs. Evidence indicates that in general 3’UTRs might function as scaffolds of different protein and RNA complexes to provide distinct functions. Diversification of 3’UTR sequences and functions inducing changes at the subcellular and cellular levels as we report in this study, also expands the possible outcomes, generating a new scale of complexity in the cell. Studying the mechanism of action of these regions intertwined with the regulation of the expression of the different variants in normal conditions and disease will facilitate the understanding of the actual role of the noncoding transcriptome in the homeostasis of the cell and the organism.

## Methods

### RNA extraction from human tissue

RNA from frozen postmortem human tissue of lumbar spinal cords of ALS patients and neuropathologically normal controls was isolated using TRIzol® reagent. All samples were obtained using ethics research protocols approved by the Western University Health Sciences Research Ethics Board (HSREB) (HSREB #103735). Control (n = 4) and ALS (n = 6) were obtained between January and September 2018. Informed consent for autopsy and retention of tissues for research purposes was obtained in written format from all patients antemortem or, if not possible, from their spouse postmortem. No minors participated in this study. In order to ensure anonymization, at the time of utilization for this research project, all cases were assigned a unique identifier with the master list correlating the unique identifier maintained in a locked cabinet in the office of the principle investigator (MJS) as per HSREB requirements. RNA quantification was performed using the NanoDrop (Thermo Fisher Scientific) and integrity was analyzed using 1 ml (50-500 ng/ml) of sample on the Agilent 2100 Bioanalyzer (Agilent Technologies Inc.) and the RNA 6000 Nano kit (Caliper Life Sciences).

### RNA-seq

Samples were sequenced at the London Regional Genomics Centre (Robarts Research Institute, London, Ontario, Canada; http://www.lrgc.ca) using the Illumina NextSeq 500 (Illumina Inc.). They were processed using the Epicentre Script-Seq v2 with Ribo-Zero rRNA reduction (Illumina Inc.). Briefly, rRNA was depleted, samples were fragmented, cDNA was synthesized, amplified and indexed by PCR. Libraries were then equimolar pooled into one library. Size distribution was assessed on an Agilent High Sensitivity DNA Bioanalyzer chip and quantitated using the Qubit 2.0 Fluorimeter (Thermo Fisher Scientific).

The library was sequenced on an Illumina NextSeq 500 as 2×76 bp paired-end runs, using two High Output v2 kits (150 cycles). Fastq data files were analyzed using Partek Flow. After importation, data was aligned to the *Homo sapiens* genome using STAR 2.5.3a and annotated using hg38 Ensembl Transcripts release 93, followed by normalization by Counts Per Million (CPM plus 0.0001).

### Alternative polyadenylation analysis (APA)

Quantitative alternative polyadenylation analysis (QAPA) (16) was performed by Quick Biology Inc. QAPA was applied using a human 3′UTR library (hg19). Briefly, salmon DNA was used to calculate the transcripts per million (TPM) value for the expression quantification of 3′UTR isoforms and QAPA was used to calculate the relative proportion of each isoform in a gene, measured as Poly(A) Usage (PAU). 3′UTRs that had a total gene expression of at least 1 TPM in at least 8/10 samples were kept in the analysis. The change in poly(A) site usage (DPAU) was defined as the difference between the median PAU of the ALS group replicates and the median PAU of the control group replicates.

### Clonings

3′UTRs of *NEFH*, *SOD1,* and *SQSTM1* detected by APA were amplified using Phusion® High-Fidelity DNA Polymerase (Thermo Fisher Scientific). The amplified fragments were initially cloned into the pGEM®-T Easy vector for sequencing and validation. Then, the 3’UTR variants were subcloned into the pEGFP-C1 expression vector for subsequent experiments.

### Cell culture and transfection

SH-SY5Y cells were cultured in Dulbecco’s Modified Eagle’s Media (DMEM) containing 10% Fetal Bovine Serum (FBS), inside an incubator at 37°C and 5% CO_2_. Cell differentiation was performed for 7 days in Eagle’s Minimum Essential Medium (EMEM) containing 2.5% FBS, 2 mM glutamine and 10 mM all-*trans* retinoic acid (Sigma-Aldrich). Transfection of eGFP-3’UTR plasmids was accomplished using Lipofectamine 2000 (Invitrogen) following the manufacturer’s instructions.

### Single-molecule FISH (smFISH, Stellaris)

SH-SY5Y cells were grown and differentiated in 6-well plates with coverslips. Fourteen hours after transfection, cells were fixed in 4% paraformaldehyde (PFA) and smFISH performed following the manufacturer’s instructions. Cells were incubated with hybridization buffer containing eGFP FISH probe labelled with CAL Fluor RED 610 dye (Stellaris, Biosearch Technologies, #VSMF-1013-5) for 4 hours at 37 °C. Nuclear staining was performed with Hoechst 32258 and coverslips were mounted using Prolong Diamond (Invitrogen). Cells were examined using an SP8 Lightening Confocal microscopy system (Leica Microsystems Inc.) and images were visualized using the LAS X 2.0 software.

### Imaging quantitative analyses

Images were processed and analyzed using Huygens Essential software (Scientific Volume Imaging) and NeuronJ plugin for ImageJ. For eGFP-*NEFH* and eGFP-*SQSTM1* 3’UTRs, deconvolution was performed using Huygens Classic Maximum Likelihood Estimation (CMLE) algorithm and cluster number and size quantification using Object Analyzer module of Huygens Essential software (Scientific Volume Imaging). For eGFP-*SOD1* 3’UTRs, filopodia were traced using NeuronJ ensuring precise segmentation. 3’UTR mRNA content within filopodia was quantified using the Huygens Object Analyzer module. Lamellipodia were quantified using Fiji (ImageJ) software by manually selecting regions of interest (ROI). The analysis was performed in 50 cells per group. Quantitative colocalization analysis was performed with Huygens colocalization module.

### Statistical analysis

A normality test was conducted for all experimental groups. When assumptions of normality or equal variances were not met, the nonparametric Kruskal-Wallis test was applied, followed by original False Discovery Rate (FDR) correction. Data distributions were visualized with violin plots to display group variability. Statistical significance was set at p<0.05 and all analyses were performed using GraphPad Prism (version 10.4.0 (621)).

## Acknowledgements

The authors gratefully acknowledge Dr. Cristian A. Droppelmann for technical assistance. This work was supported by The McFeat Family Fund, the Michael Hall Foundation and the Canadian Institutes of Health Research (CIHR) grant SOP-160442.

## Abbreviations

3′UTR: 3′ Untranslated Region
ALS: Amyotrophic Lateral Sclerosis
APA: Alternative Polyadenylation
CMLE: Classic Maximum Likelihood Estimation
*NEFH*: Neurofilament Heavy Chain
PAP: Poly(A) Polymerase
PAS: Polyadenylation Signal
PAU: Polyadenylation Usage
QAPA: Quantitative Alternative Polyadenylation Analysis
*SOD1*: Superoxide Dismutase 1
*SQSTM1*: Sequestosome 1
smFISH: Single-Molecule Fluorescence *in situ* Hybridization
TPM: Transcripts Per Million.

## Supporting information

**S1 Fig. RNA-seq analyzed using QAPA from ALS human spinal cords.** (A) PCA plot of RNAseq/QAPA that shows distribution of ALS (A) and control (C) spinal cord samples. (B) Heat map of poly(A) site usage (PAU) of the proximal 3’UTR isoform (DDPAU).

**S2 Fig. *SQSTM1* 3**′**UTR variants doesn’t modify RNA cluster or filopodia formation.** RNA-FISH was performed in differentiated SH-SY5Y cells transfected with eGFP-alone (Control), *SQSTM1* 3′UTR-Short, Medium or Long. (A) Schematic representation of the *SQSTM1* 3′UTR variants. (B) The expression of *SQSTM1* 3’UTRs show no alterations in RNA cluster formation. (C) The expression of *SQSTM1* 3’UTRs shows no changes in filopodia formation. Scale bar, 5 μm.

**S3 Fig. Quantification of cluster areas for *NEFH* 3**′**UTR variants.** RNA-FISH was performed in differentiated SH-SY5Y cells transfected with eGFP-alone (Control), NEFH 3′UTR-Short, or Long (red). Violin plots depicting the area of RNA clusters smaller than 1.3 μm^2^ in the nucleus and the cytoplasm. Statistical comparisons were conducted using a nonparametric Kruskal–Wallis test followed by original FDR correction. P-values are indicated above each comparison.

**S4 Fig. *SOD1* 3′UTR variants affect lamellipodia formation.** (A) RNA-FISH performed in differentiated SH-SY5Y cells transfected with *SOD1* 3′UTR-Short, Medium, or Long constructs. Representative confocal images of lamellipodia are highlighted with squares. Scale bar, 10 mm. (B) Violin plots showing number of lamellipodia and lamellipodia area. Statistical significance was determined by nonparametric Kruskal–Wallis test followed by original false discovery rate (FDR) correction, with p-values indicated above each comparison.

**S5 Fig. *SOD1* 3**′**UTR variants colocalize with F-actin.** RNA-FISH was performed in differentiated SH-SY5Y cells transfected with *SOD1* 3′UTR-Short, Medium, or Long, followed by immunofluorescence to detect F-actin. *SOD1* 3′UTR (red) and F-actin (green) colocalize in the cytoplasm and neurites. Overlap is observed in yellow. The resulting overlap values (R) for each region are indicated in each image. Based on R values for cytosol and neurites, moderate colocalization is observed for *SOD1 3*′UTR-Short, high colocalization for *SOD1* 3′UTR-Medium and high colocalization for *SOD1* 3′UTR-Long. Scale bar, 10 μm.

