## Supplementary figures and images for "3’UTR variants of ALS-linked RNAs modify subcellular and cellular phenotypes"

### Supplemental figure 1

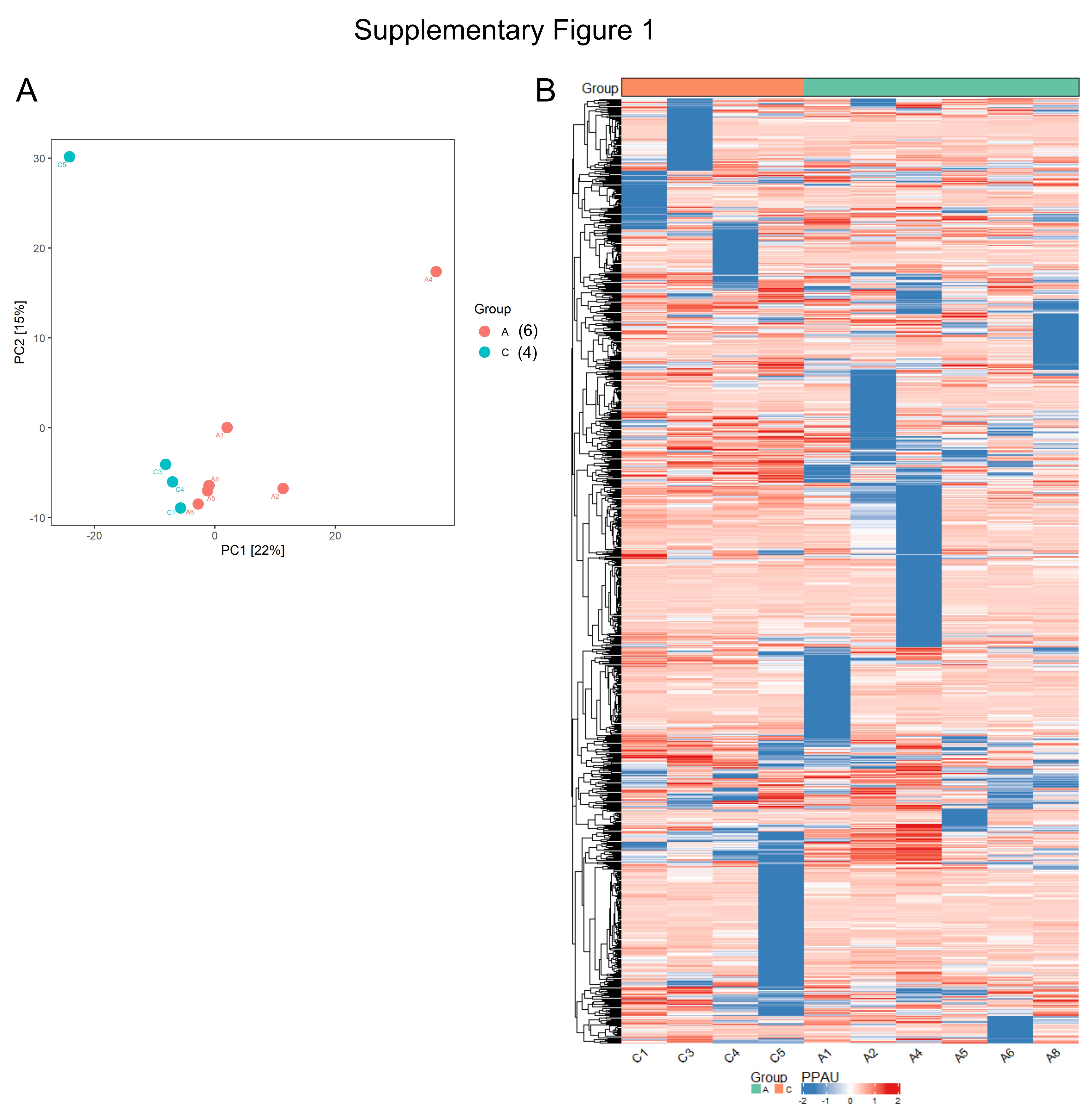

### Supplemental figure 2

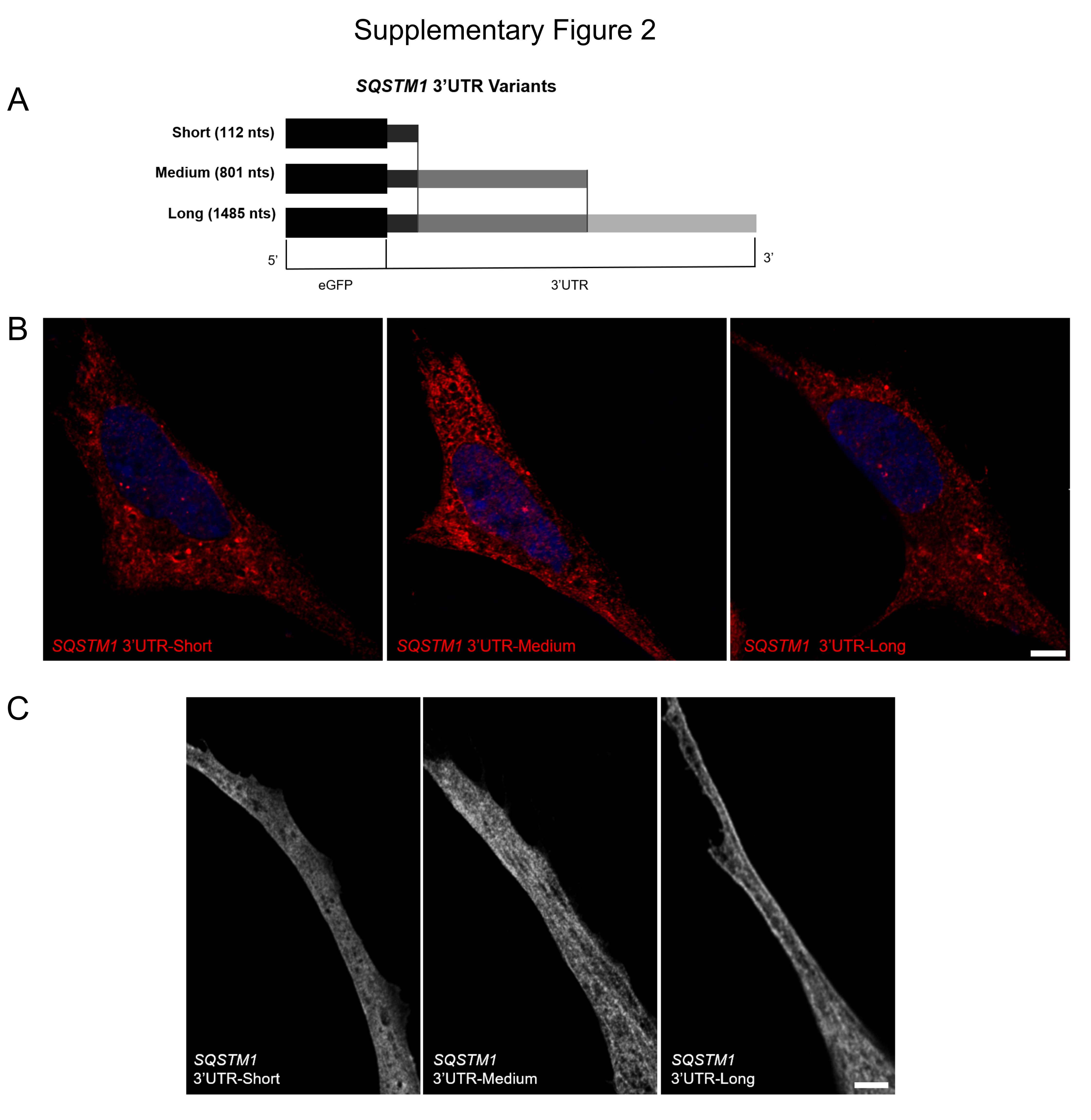

### Supplemental figure 3

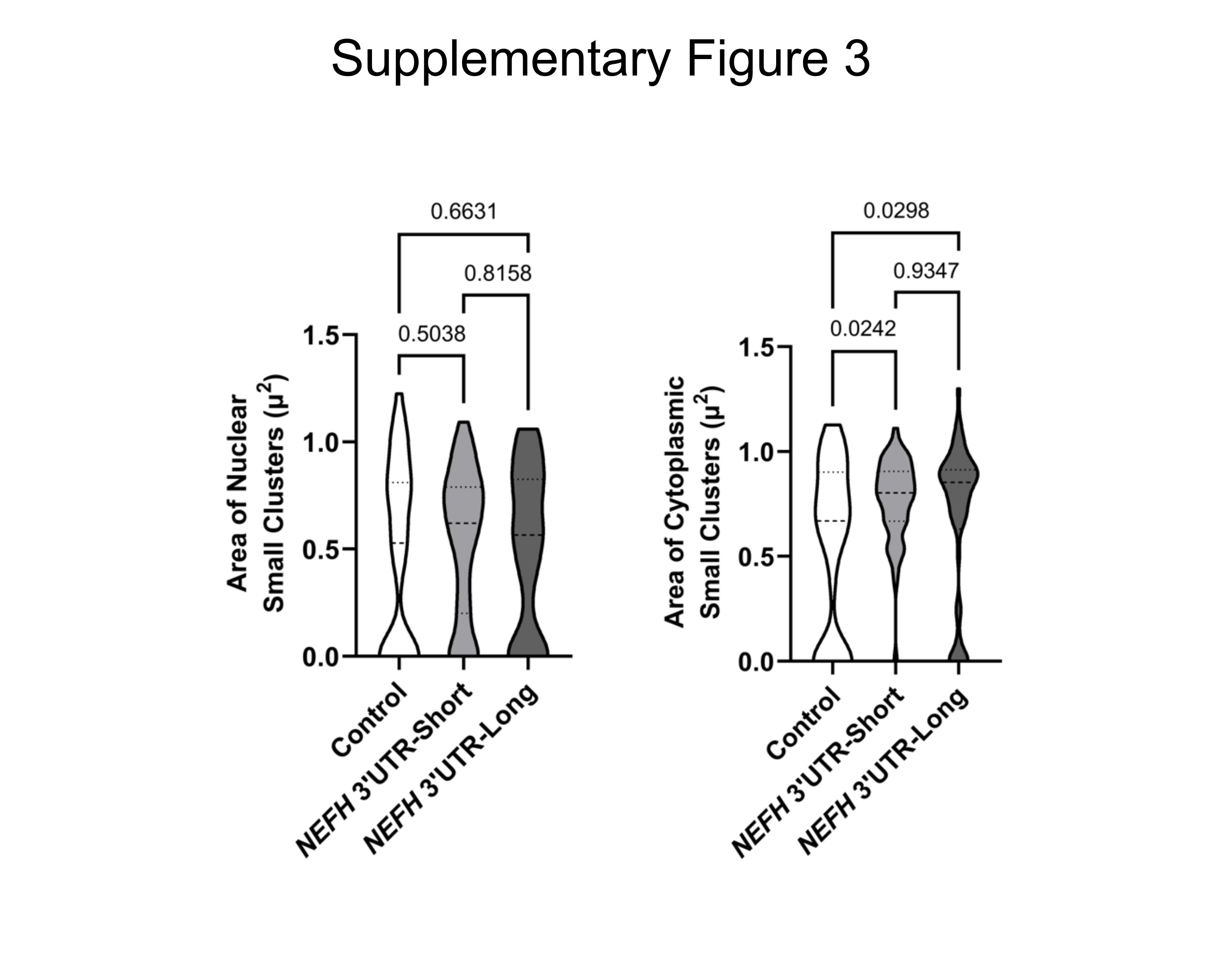

### Supplemental figure 4

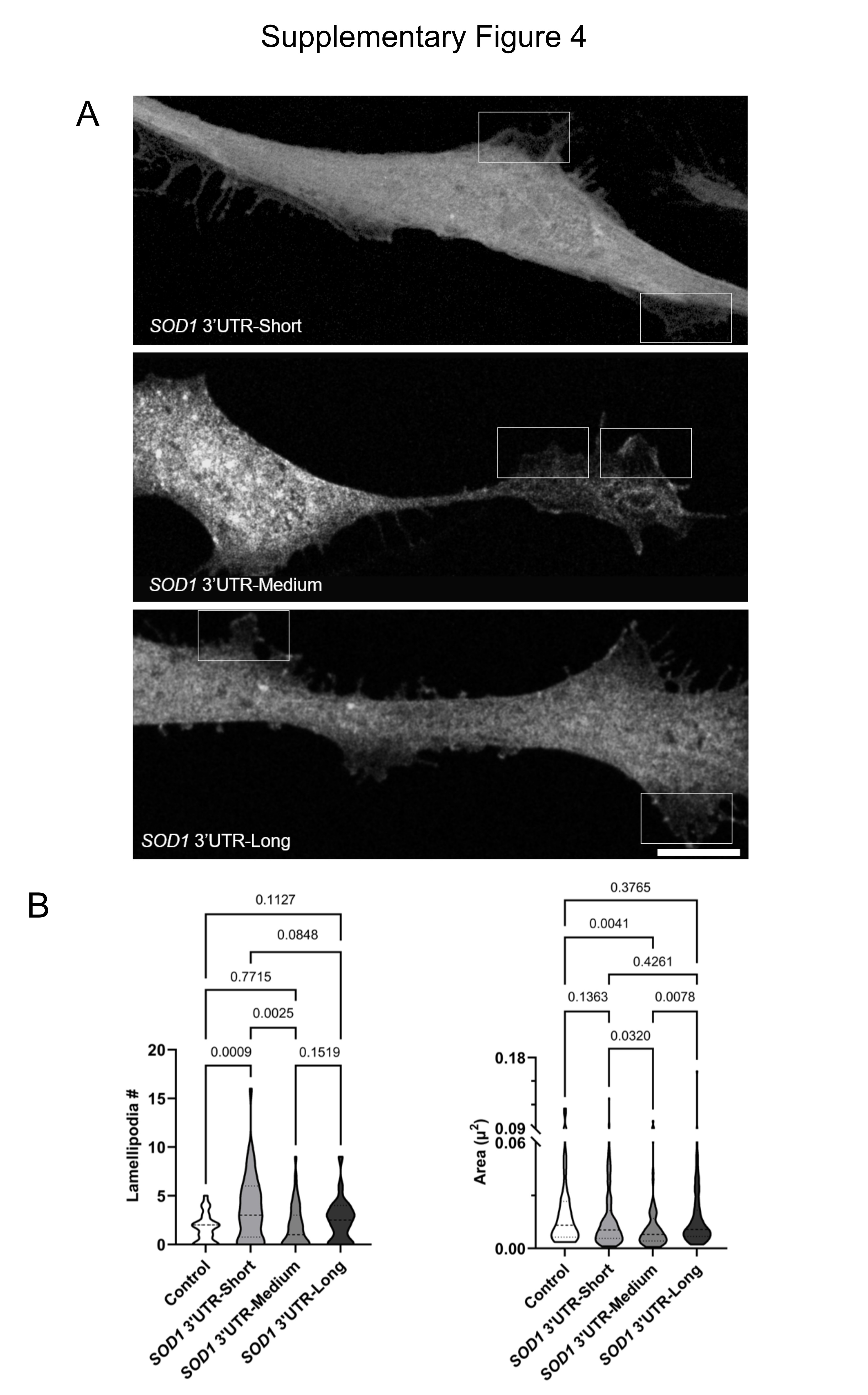

### Supplemental figure 5

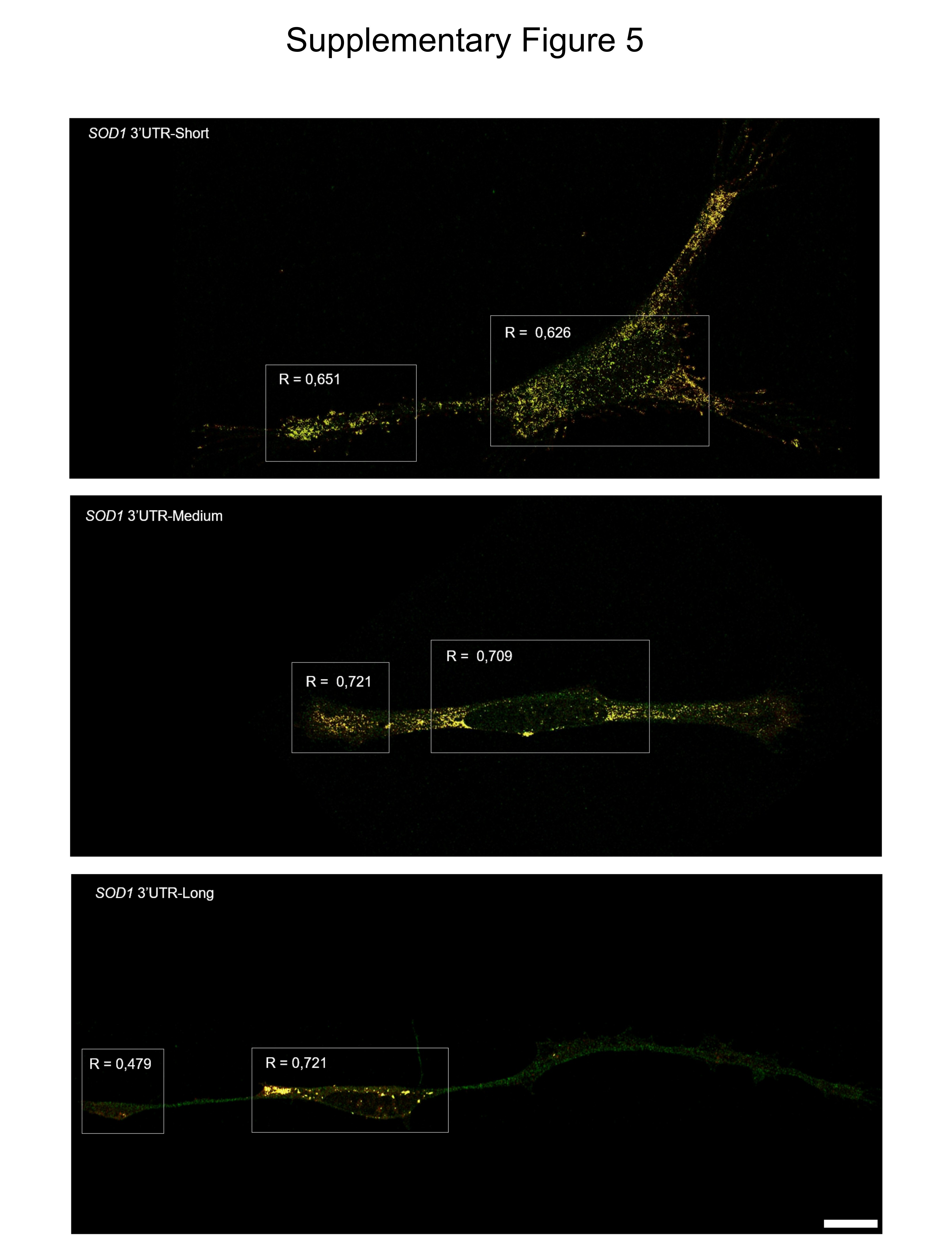
